# Serology reflects a decline in the prevalence of trachoma in two regions of The Gambia

**DOI:** 10.1101/149237

**Authors:** Stephanie J Migchelsen, Nuno Sepúlveda, Diana L Martin, Gretchen Cooley, Sarah Gwyn, Harry Pickering, Hassan Joof, Pateh Makalo, Robin Bailey, Sarah E. Burr, David CW Mabey, Anthony W Solomon, Chrissy h Roberts

## Abstract

Trachoma is caused by *Chlamydia trachomatis* (Ct). It is targeted for global elimination as a public health problem. In 2014, a population-based cross-sectional study was performed in two previously trachoma-endemic areas of The Gambia. Participants of all ages from Lower River Region (LRR) (N = 1028) and Upper River Region (URR) (N = 840) underwent examination for trachoma and had blood collected for detection of antibodies against the Ct antigen Pgp3, by ELISA. Overall, 30 (1.6%) individuals had active trachoma; the prevalence in children aged 1–9 years was 3.4% (25/742) with no statistically significant difference in prevalence between the regions. There was a significant difference in overall seroprevalence by region: 26.2% in LRR and 17.1% in URR (p<0.0001). In children 1-9 years old, seroprevalence was 4.4% in LRR and 3.9% in URR. Reversible catalytic models using information on age-specific seroprevalence demonstrated a decrease in the transmission of Ct infection in both regions, possibly reflecting the impact of improved access to water, health and sanitation as well as mass drug administration campaigns. Serological testing for antibodies to Ct antigens is potentially useful for trachoma programmes, but consideration should be given to the co-endemicity of sexually transmitted Ct infections.

## INTRODUCTION

Trachoma is caused by ocular infection with the obligate intracellular bacterium *Chlamydia trachomatis* (Ct). It is the leading infectious cause of blindness worldwide^1^. Infection is associated with clinical signs of inflammation in the conjunctiva, known as active trachoma; these include trachomatous inflammation—follicular (TF) and trachomatous inflammation—intense (TI). Many repeated episodes of active trachoma over years to decades can lead to trachomatous trichiasis (TT), which may lead to impaired vision. The World Health Organization (WHO) estimates that over 200 million people in 42 countries are at risk of blindness from trachoma ^2^. In 2010, approximately 1.9 million people suffered from visual impairment or blindness due to trachoma^1^. The WHO Alliance for the Global Elimination of Trachoma by 2020 (GET2020) aims to eliminate trachoma as a public health problem by 2020^2^ through the SAFE Strategy (Surgery, Antimicrobials, Facial cleanliness and Environmental improvement). The WHO endorsed strategy for global control is to reduce the population prevalence of TF to <5% in children aged 1–9 years and the prevalence of unmanaged TT to <0.2% in adults aged 15 years and above^3^. By 2014, seven countries had reported having met these targets nationally^4^. The Gambia has been a hub for trachoma research for over 50 years and is on course to declare the elimination of trachoma as a public health problem by 2020. Two National Surveys of Blindness and low vision in The Gambia demonstrated that there had been a nation-wide decreases in the prevalence of active trachoma and TT between 1986 and 2000^22^,^23^. Specific elimination efforts, undertaken between 2007 and 2010 and run by the National Eye Health Programme (NEHP), included mass drug administration (MDA) of azithromycin in 23 districts across the country. The Partnership for the Rapid Elimination of Trachoma (PRET)^5^ was embedded within the national programme and measured the prevalence of Ct infection in children residing in four districts in which MDA had been administered. Deployment of interventions and disease control measures across The Gambia was not even, with some districts receiving the full SAFE intervention and others receiving reduced (F & E components) or no specifically targeted Interventions. It is important to be able to evaluate the effectiveness of these interventions and a significantly reduced or halted transmission of infection is a key indicator that elimination may have been achieved.

The current guidelines for post-intervention surveillance of Ct transmission intensity are based on the prevalence of TF in children aged 1–9 years. This is problematic because in the peri-elimination and low-endemicity settings^5^,^6^ the correlation between the clinical signs of trachoma and the presence of ocular Ct infection is poor. This diminishment of the positive predictive value of clinical signs means that the false positive rate of TF screening is increased and the specificity of the clinical sign is negatively affected.

Reduced global endemicity also leads to the unmasking of diseases that resemble trachoma clinically but which have no clear link to Ct infections. We recently surveyed the western division of the Solomon Islands^7^, a population where TF was not found to be a highly specific indicator of ocular Ct infection. Around 26% of 1-9 year olds living there had TF, but Ct infection was very scarce at just 1.3%. Using a serological tool we showed that 53/66 (80%) of the cases of TF that we observed were in people who were serologically negative for prior Ct infection^8^. Clinical signs of trachoma were also a poor indicator for the need to deploy antimicrobials in Fiji, where the high prevalence of TT cases could be better explained by socio-epidemiological practices of eyelash depilation than by Ct infections^9^. The current WHO guidelines would recommend MDA in both Fiji and the Solomon Islands, but the evidence from more detailed surveys makes an argument against the likelihood that the use of antimicrobials would be effective. Situations such as this highlight the need to develop new tools that can support programmes to make informed decisions about how to use antimicrobials in trachoma control. Antibodies against Chlamydia trachomatis reflect cumulative exposure to Ct^10^,^11^ and it has been suggested that programmes could use some measure of seroprevalence as an alternative indicator of changes in transmission^12^,^13^. Previous work has investigated the use of age-specific seroprevalence for surveillance in the peri-^14^ and post-MDA setting^13^,^15^,^16^. Serological techniques for the detection of antibodies against Ct have been used to study the epidemiology of both urogenital and ocular Ct infections^17^,^18^. An enzyme-linked immunoassay (ELISA) which detects antibodies against Pgp3 (pCT03) has recently been used for analysis of samples collected from trachoma-endemic regions^19^,^20^. The Pgp3 molecule is an immunogenic Ct-specific protein that is encoded by the Ct plasmid and which is highly conserved at the genetic level among Ct isolates^21^.

In the current study we have used serological data from a Pgp3-specific ELISA to gain insight into the dynamics of Ct transmission in two regions of The Gambia, one of which (the Lower River Region [LRR]) had received three annual rounds of MDA, whilst the other (the Upper River Region [URR]) had not. URR was not targeted for antimicrobial use because the TF prevalence had already dropped below the WHO threshold for elimination as a public health problem.

To explore the utility of serology as a tool for surveillance in trachoma elimination programmes, we have modelled the seroconversion rate (SCR) in URR and LRR under different epidemiological settings. The SCR is the yearly average rate by which seronegative individuals become seropositive because of disease exposure. SCR is a metric that acts as a surrogate measure for the underlying force of infection (FoI) and the age specific SCR can be used to model changes in FOI across time, potentially identifying periods in which there were substantial changes in the rate of transmission of Ct infection in the population.

## RESULTS

We recruited participants of all ages from LRR (n=1028, 41.9% male) and URR (n=840, 42.5% male). Ten participants were excluded from the study because they either declined to provide a blood sample (n=1) or had incomplete examination data (n=9). The median participant age was 13 years in LRR (range: 1-88, IQR 6–34) and 11 years in URR (range: 0–90, IQR 5–40). The proportion of participants by age group is shown in Table 1. There were significantly more females than males overall (X^2^=45.332, p<0.0001), which held true in both LRR (X^2^=26.483, p<0.0001) and URR (X^2^=18.601, p<0.0001) and is representative of Gambian demographics^24^. Age distributions were approximately equal between the two regions, as seen in Table 1 (X^2^=27.703, p=0.14).

**Table 1.** Age distribution of study participants, Lower River Region and Upper River Region, The Gambia, 2014.

|  | Both Regions |  | Lower River Region |  | Upper River Region |  |
| --- | --- | --- | --- | --- | --- | --- |
|  | N | % | N | % | N | % |
| **Overall** | 1832 |  | 1010 | 55.1 | 822 | 44.9 |
| **Gender** |  |  |  |  |  |  |
| Female | 1056 | 57.7 | 584 | 58.0 | 472 | 57.0 |
| Male | 776 | 42.2 | 426 | 42.0 | 350 | 43.0 |
| **Age group (years)** |  |  |  |  |  |  |
| 1-9 | 742 | 39.7 | 383 | 37.3 | 359 | 42.7 |
| 10-19 | 412 | 22.1 | 231 | 22.5 | 181 | 21.5 |
| 20-29 | 191 | 10.2 | 101 | 9.8 | 90 | 10.7 |
| 30-39 | 152 | 8.1 | 79 | 7.7 | 73 | 8.7 |
| 40+ | 335 | 17.9 | 216 | 21.0 | 119 | 14.2 |

### Clinical signs

Thirty cases of TF were found in total (1.6%, 95% CI=1.1–2.3%), of which 25 were in children aged 1– 9 years (3.4%, 95% CI=2.2–4.9, Table 2). There were 78 cases of trachomatous conjunctival scarring (TS), of which 70 were in participants aged 15 years and above (8.3%, 95%CI=6.6–10.5); eight cases of TT, all of which were in those aged 40 years and older (0.9%, 95% CI=0.4–1.8); and one case of corneal opacity (CO), in a participant over 40 years of age (0.1%, 95% CI=0.0–0.6) (Table 2). The prevalence of TS was significantly different (X^2^=7.7932, p=0.005) between LRR (5.4%, 95% CI=4.1-7.1%) and URR (2.8%, 95% CI=1.8–4.2%). The TF prevalence in children aged 1–9 years was not significantly different (X^2^=0.199, p=0.66) between LRR (3.7%, 95% CI=2.1-6.2%) and URR (3.1%, 95%CI=1.6-5.6%). The prevalence of TI, TT and CO in this population was too low for further statistical analysis.

**Table 2.** Frequency of signs of trachoma in study participants, Lower River Region and Upper River Region, The Gambia, 2014.

|  | Frequency of signs (%) |  |  |  |  |  |
| --- | --- | --- | --- | --- | --- | --- |
|  | N | TF | TI | TS | TT | CO |
| **Overall** | 1832 | 30 (1.6) | 4 (0.2) | 78 (4.3) | 8 (0.4) | 1 (0.1) |
| **Region** |  |  |  |  |  |  |
| LRR | 1010 | 18 (1.8) | 4 (0.4) | 55 (5.4) | 7 (0.7) | 1 (0.1) |
| URR | 822 | 12 (1.4) | 0 | 23 (2.8) | 1 (0.1) | 0 |
| **Gender** |  |  |  |  |  |  |
| Female | 1056 | 10 (0.9) | 3 (0.3) | 52 (4.9) | 5 (0.5) | 1 (0.1) |
| Male | 776 | 20 (2.5) | 1 (0.1) | 26 (3.4) | 3 (0.4) | 0 |
| **Age group (years)** |  |  |  |  |  |  |
| 1–9 | 742 | 25 (3.4) | 2 (0.3) | 7 (1.1) | 0 | 0 |
| 10–19 | 412 | 4 (1.0) | 1 (0.2) | 2 (0.5) | 0 | 0 |
| 20–29 | 191 | 0 | 0 | 1 (0.5) | 0 | 0 |
| 30–39 | 152 | 1 (0.7) | 1 (0.7) | 5 (3.3) | 0 | 0 |
| 40+ | 335 | 0 | 0 | 63 (18.8) | 8 (2.4) | 1 (0.3) |
| 1–9 year olds -LRR | 383 | 14 (3.7) | 2 (0.5) | 1 (0.3) | 0 | 0 |
| 1–9 year olds -URR | 359 | 11 (3.1) | 0 | 6 (1.7) | 0 | 0 |
| ≥10 year olds-LRR | 645 | 4 (0.6) | 2 (0.3) | 54 (8.4) | 7 (1.1) | 1 (0.2) |
| ≥10 year olds-URR | 481 | 1 (0.2) | 0 | 17 (3.5) | 1 (0.2) | 0 |

TF=trachomatous inflammation—follicular; TI=trachomatous inflammation—intense;
TS=trachomatous conjunctival scarring; TT=trachomatous trichiasis; CO=corneal opacity
LRR=Lower River Region; URR=Upper River Region

Data from this study show very low prevalence of active trachoma in these regions. In children aged 1–9 years, only 4.2% had detectable antibodies against Pgp3. In both regions, the prevalence of TF in 1–9-year-olds was below the 5% threshold for elimination as a public health problem as specified by the WHO.

### Antibody responses in the populations

The threshold for seropositivity was set using a finite mixture model^25^ to classify the samples as seropositive or seronegative based on maximum likelihood methods. The threshold was set at the mean of the Gaussian distribution of the seronegative population plus four standard deviations, 0.810 OD_450nm_ to ensure high specificity. Previous studies using the Pgp3 ELISA have commonly used a threshold set as three standard deviations above the mean of the negative population (the 97.5% confidence interval) ^19^,^26^. Using that lower threshold resulted in the same qualitative conclusions being drawn (See Supplementary Information).

The seroprevalence of antibodies against Pgp3 for each region, by age and gender, is summarised in Table 3. The overall seroprevalence in LRR and URR was 26.2% (95% CI=23.5–29.0) and 17.1% (95% CI=14.7–19.9%), respectively. Adjusting for multiple comparisons, there was a significant difference in overall seroprevalence between the two regions (X^2^=20.72, p<0.0001). Figure 1 shows the seroprevalence by age group and region. In children 1-9 years old, seroprevalence was 4.4% in LRR and 3.9% in URR. As expected, the prevalence of anti-Pgp3 antibodies increased with age using the non-parametric test for trend (z-score=23.35, p<0.0001; alpha=0.01). The seroprevalence doubled between 10–19-year-olds and the next oldest age group, 20–29-year-olds, both in the two regions combined, and in each region. Across both regions, in study participants who had no signs of trachoma, seroprevalence was 21% (95% CI = 19–23). Of those who had active trachoma (TF and/or TI) (n=30), 3% were seropositive (95% CI = 0.1–17) and of those who had scarring trachoma (TS and/or TT and/or CO in either eye; n=122) 56% were seropositive (95% CI = 44–67).

**Figure 1.**
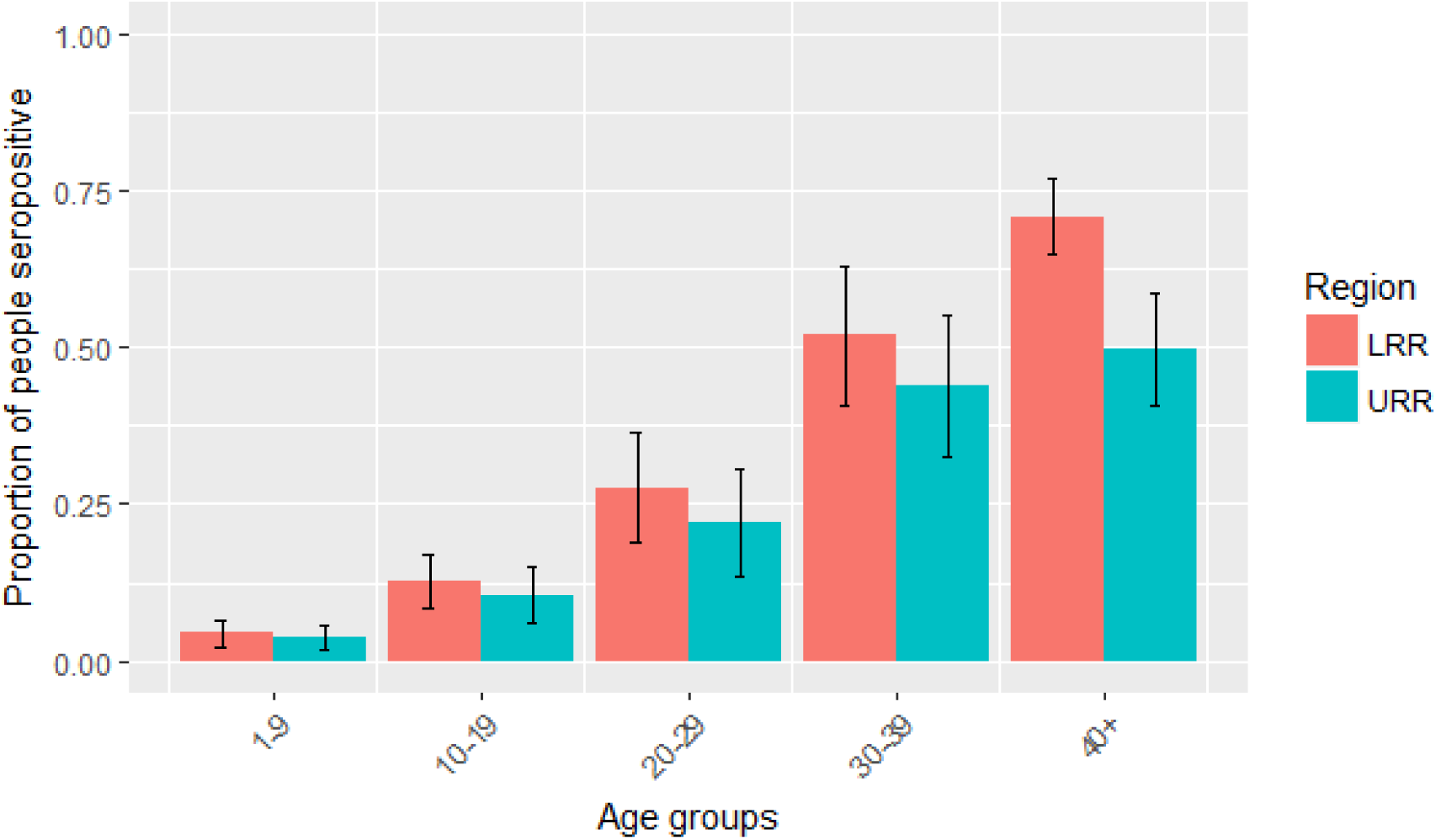
Proportion of participants who were seropositive for anti-Pgp3 antibodies by age group and region, Lower River Region (LRR) and Upper River Region (URR), The Gambia, 2014. Vertical bars indicate 95% CIs.

**Table 3.** Seroprevalence of anti-Pgp3 antibodies by region, gender and age, Lower River Region and Upper River Region, The Gambia, 2014.

|  | **Both** |  |  |  | **Lower River Region** |  |  |  | **Upper River Region** |  |  |  |
| --- | --- | --- | --- | --- | --- | --- | --- | --- | --- | --- | --- | --- |
|  | **N** | **n** | **%** | **95% CI** | **N** | **n** | **%** | **95% CI** | **N** | **n** | **%** | **95% CI** |
| **Overall** | 1832 | 412 | 22.5 | 20.2-24.0 | 1010 | 268 | 26.5 | 23.5-29.0 | 822 | 144 | 17.5 | 14.7-19.9 |
| **Gender** |  |  |  |  |  |  |  |  |  |  |  |  |
| Female | 1056 | 295 | 27.9 | 24.8-30.2 | 584 | 190 | 32.5 | 28.8-36.5 | 472 | 105 | 22.2 | 18.6-26.3 |
| Male | 776 | 117 | 15.1 | 12.4-17.5 | 426 | 78 | 18.3 | 14.8-22.4 | 350 | 39 | 11.1 | 8.1-15.0 |
| **Age group (years)** |  |  |  |  |  |  |  |  |  |  |  |  |
| 1–9 | 742 | 31 | 4.2 | 2.9-5.9 | 383 | 17 | 4.4 | 2.6-7.0 | 359 | 14 | 3.9 | 2.1-6.5 |
| 10–19 | 412 | 48 | 11.7 | 8.7-15.1 | 231 | 29 | 12.6 | 8.6-17.5 | 181 | 19 | 10.5 | 6.4-16.0 |
| 20–29 | 191 | 48 | 25.1 | 19.1-31.9 | 101 | 28 | 27.7 | 19.3-37.5 | 90 | 20 | 22.2 | 14.1-32.2 |
| 30–39 | 152 | 73 | 48.0 | 39.9-56.3 | 79 | 41 | 51.9 | 40.4-63.3 | 73 | 32 | 43.8 | 32.2-55.9 |
| 40+ | 335 | 212 | 63.3 | 57.9-68.5 | 216 | 153 | 70.8 | 64.3-76.8 | 119 | 59 | 49.6 | 40.3-58.9 |

The prevalence of antibodies was significantly higher in females than in males (z-score=6.384, p<0.0001). The same was true in each region (LRR: z-score=4.881, p<0.0001; URR: z-score=4.114, p<0.0001). When data were considered for each age group, the seropositivity difference between males and females was only significant for 10–19-year-olds (z-score=2.667, p=0.0077) and 30–39-year-olds (z-score=0.2551, p=0.0107). Seroprevalence by gender and age group is shown in Figure 2.

**Figure 2.**
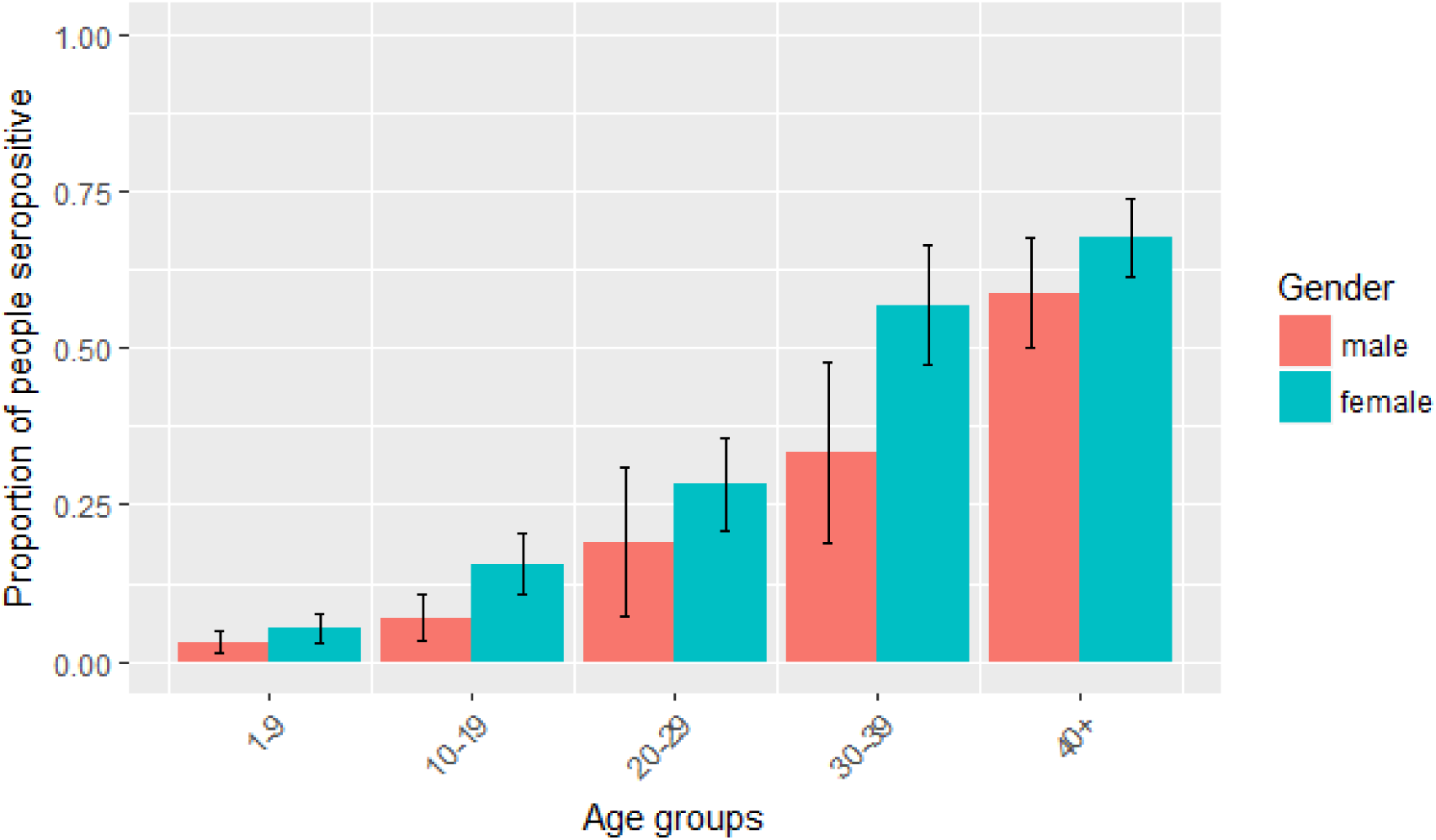
Proportion of participants who were seropositive for anti-Pgp3 antibodies, by age group and gender, Lower River Region and Upper River Region, The Gambia, 2014. Vertical bars indicate 95% confidence intervals.

### Reduction in seroconversion rates over time

We used a reversible catalytic model together with the profile likelihood method to identify reductions in seroconversion rates (SCR) in both LRR and URR. The reversible catalytic model is based on the premise that individuals transit between seropositive and seronegative states with specific average rates^27^. We compared three different models under which transmission dynamics might have changed across time. Our first model used the assumption that there had been a constant transmission, the second that there had at some point been an abrupt change in force of infection and the third that there had been a constant FOI to a point, after which it had decayed in a log-linear manner. We found that the model of abrupt change was most likely to explain the data we had observed in The Gambia. Abrupt reductions in SCRs were identified in both LRR and URR (Table 5 and Figure 3) and these changes in SCR are estimated to have occurred respectivekty 23 and 16 years before data collection took place (Figure 3). For LRR, the SCR would appear to have dropped from an incidence of 0.062 yearly events per person, to 0.010 yearly events per person. These estimates implied a putative 6.3-fold decrease in transmission intensity. In URR, the estimate of the past SCR dropped from 0.050 yearly events per person to 0.008 yearly events per person, a 3.2-fold decrease (Table 5). Figure 4 shows the expected seroprevalence curves as function of age assuming a change in transmission intensity.

**Figure 3.**
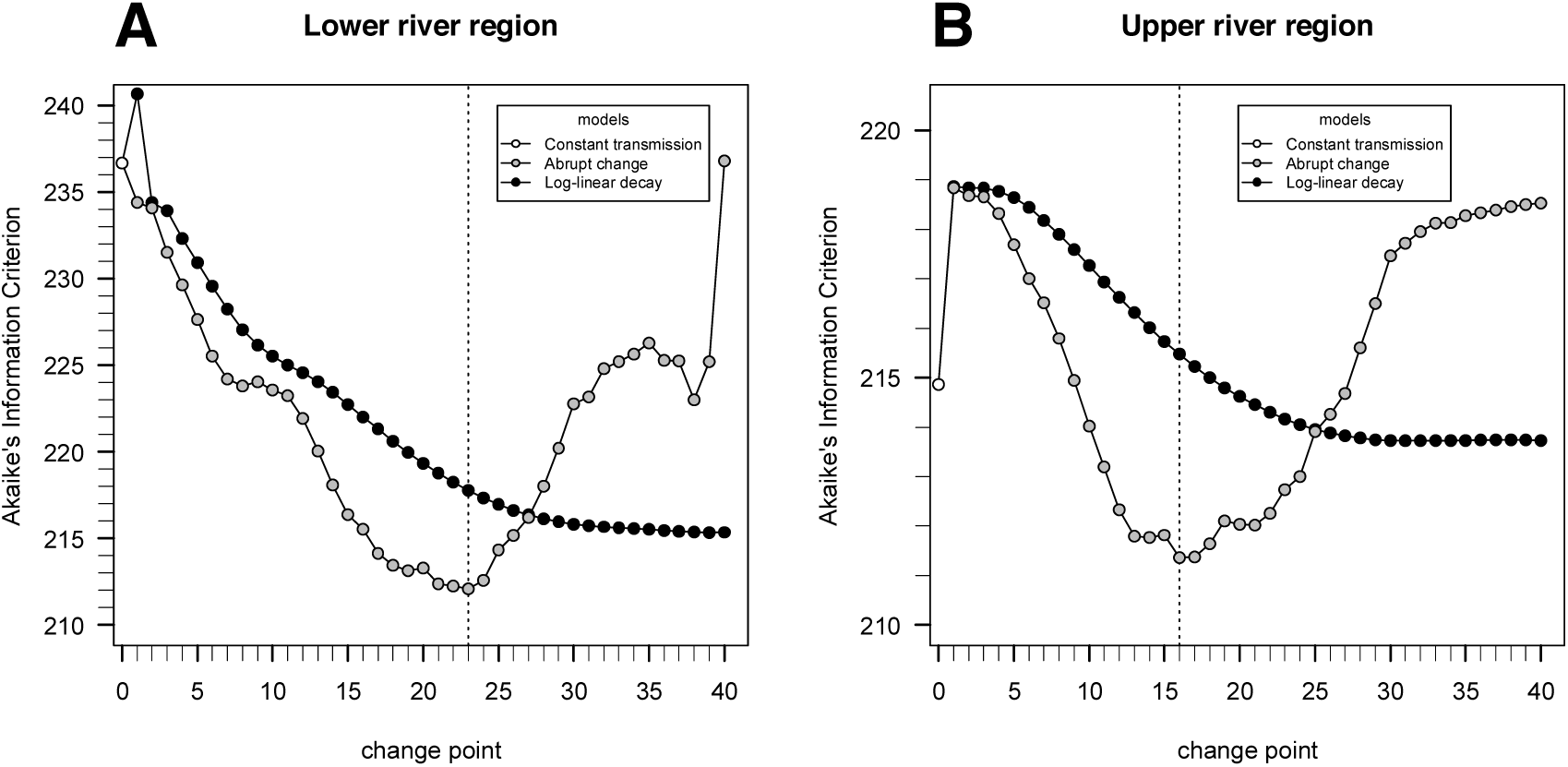
Akaike’s information criterion (AIC) using the profile likelihood method for estimating the change-point for the models assuming an abrupt reduction in transmission intensity or annual log-linear decay of transmission intensity from the change-point to the present. In this analysis, the best model as function of change point is the one that leads to the minimum estimate of the AIC. Note that a change point of 0 is equivalent to the simple model assuming a constant transmission intensity over time. The results suggest abrupt reductions of transmission intensity 23 and 16 years before sampling for respectively the lower and upper river regions.

**Table 5.** Maximum likelihood estimates for the past and current seroconversion and seroreversion rates (SCR and SRR, respectively) associated with data collected from participants in Lower River Region and Upper River Region, The Gambia, 2014 where the respective 95% confidence intervals are shown in brackets.

| Region | SCR<sub>past</sub> | SCR<sub>current</sub> | SRR | Fold change |
| --- | --- | --- | --- | --- |
| Lower River Region | 0.095<br>(0.051-0.176) | 0.015<br>(0.012-0.019) | 0.008<br>(0.004-0.015) | 6.3 |
| Upper River Region | 0.038<br>(0.018-0.082) | 0.012<br>(0.009-0.016) | 0.011<br>(0.003-0.039) | 3.2 |

## DISCUSSION

The Gambian government is currently compiling evidence for validation of trachoma elimination, following completion of its three-year TF/TT surveillance plan, which began in 2011^28,29^. Previous studies have demonstrated declines in the prevalence of active trachoma in The Gambia both prior to^6,30^ and in response to the implementation of specific trachoma-control interventions^31–33^. Increased access to water, education and healthcare in The Gambia during recent decades are thought to have had an impact^34^, manifesting in a secular decline. Regardless of cause, the prevalence of active trachoma in 0–14-year-olds fell from 10.4% in 1986^23^ to less than 5% in 1996^22^ and has subsequently remained low. Prevalence data from historical studies is presented in Table 6. More recently*, six years* before the survey reported here, communities in LRR received azithromycin to treat ocular Ct infection, further reducing the prevalence of trachoma in this region^5,29^, while communities in URR did not. Although not measured in our study, previous work has shown that there has been a reduction in the prevalence of ocular Ct infection in two villages in LRR^35,36^ with the most recent measurements showing 0.5% infection prevalence in PRET villages, a portion of which are in LRR^5^. In line with the findings of those previous studies, we provide a further data point showing TF prevalence <5% for each region. The prevalence of clinical signs strongly supports The Gambia’s claim to have eliminated trachoma as a public health problem from these areas. The seroprevalence estimates and FoI modelling of trachoma transmission intensity offer further support that transmission is greatly reduced and that trachoma is no longer a public health problem in the areas studied. Seroprevalence data presented here are very similar to those seen in a recent study in Tanzania, where ocular Ct infection was eliminated in 2005^37^ and serology shows an equally low prevalence of antibodies against Ct in children^13^.

**Table 6.**
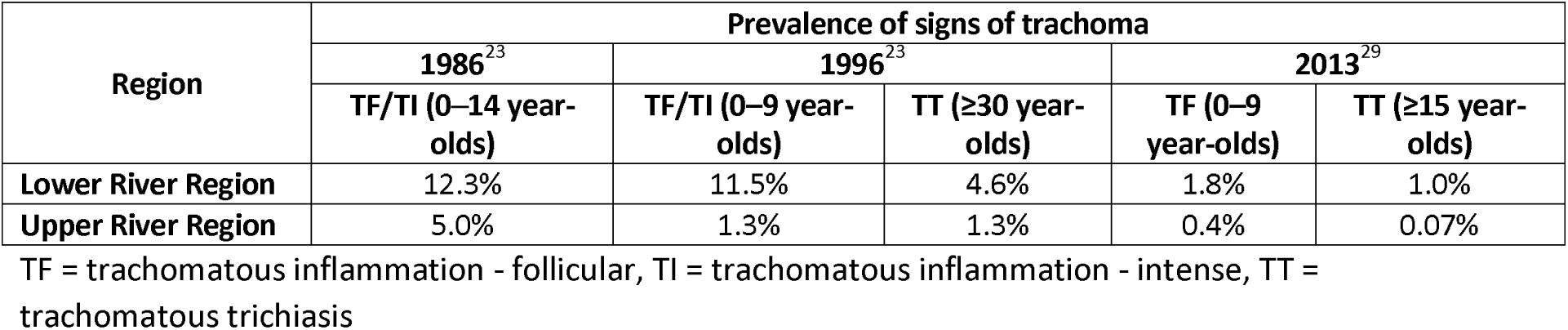
Previously published data on the prevalence active trachoma and trachomatous trichiasis, Lower River Region and Upper River Region, The Gambia, 1986–2013. Data from the 1986 survey was not available for 0–9 year olds, thus we have used the data for 0–14 year olds.

We used three SCR models to estimate the change in SCR: the first model assumed no change in SCR and serves as a baseline for the second and third models, both of which did assume a change in SCR. The best model allowed for an acute change in SCR (Table 5, Figure 3).

Our analysis of age-specific seroprevalence suggests that the FOI is currently very low, with a substantial decrease in SCR in children having occurred approximately 20 years ago (compared to the children that grew up before then). This change is too acute to reflect secular decline in the FoI of trachoma and may indicate the effects of interventions.

One confounding factor in this analysis is that the serological test we used was not specific to ocular Ct infections, rather it indicates whether the individual being tested has antibodies against Ct that might originate from historical ocular or urogenital infections. Whilst the acute change in SCR that we identified would be consistent with a significant drop in the transmission intensity of ocular Ct in the mid to late 1990s, it could also be explained by a confounding signal from contemporary seroconversion events that relate to sexual activity of people in their late teens to mid-twenties. It is arguably most likely that the data reflect both things, but without current data describing the population prevalence of urogenital and ocular infections, the proportional contributions of STIs and trachoma to the change in SCR cannot be fully assessed.

No currently available serological test distinguishes between exposure to ocular and urogenital CT infection. It is interesting to note that the seroprevalence among 10 - 19-year-old Gambian females in our study was almost double that of their male counterparts of the same age. In the Gambia, the median age at which females first have sex is 18.6 years, with 52% of women aged 20 - 24 years surveyed as part of the Demographic and Health Survey (2013) having had sexual intercourse by age 20. In males the median age is 23.1 years, with 48% of men aged 25–29 years surveyed having had sexual intercourse by age 22^39^. This could in part explain why we observed a gender difference in seroprevalence in the 10 - 19-year-old age range, as those with earlier sexual debut would be expected to be more likely to acquire STIs^40–42^ and to seroconvert.

In a population in which the urogenital Ct infection prevalence has been consistently low, it might be expected that anti-Pgp3 serological data would more accurately reflect longitudinal trends in ocular Ct transmission. Whilst data suggest that the prevalence of urogenital Ct infections has historically been very low in rural areas of The Gambia^43,44^, there are no recent data, nor data based on modern molecular testing methods. A 2003 study from Malicounda in Thiès Region of neighbouring Senegal estimated that the prevalence of urogenital Ct infection there was just 0.3% (n=73)^45^. A systematic review of global estimates of incidence and prevalence of sexually transmitted infections (STI), including urogenital chlamydia, estimated that the prevalence of urogenital chlamydia was 2.9% in low-income countries^46^, although it is noted that this study did not include data from The Gambia.

The cross-specificity of serological tests for antibodies emerging in response to ocular and urogenital Ct infections is a substantial hurdle that will need to be overcome if serological tests are to be widely deployed for trachoma monitoring.

The confidence intervals associated with SCRs seen in Figure 4 are very broad (which in part reflects the uncertainty of modelling approaches) and comparison between the charts for LRR and URR is indicative but not conclusive of a difference in SCR between the two regions. Although a larger sample size would reduce the uncertainty, interpretation of the model depends to a large extent on the magnitude of the change as well as the timing between the change in SCR and sample collection, as seen in malaria modelling work^25^. The very large sample sizes required of studies that could delineate SCR changes with high precision could be prohibitive. The analysis is further limited by the fact that the model assumes a memoryless property over time and the SCR models are defined as a function of the seroreversion rate, which not easily estimated from a single cross-sectional survey.

**Figure 4.**
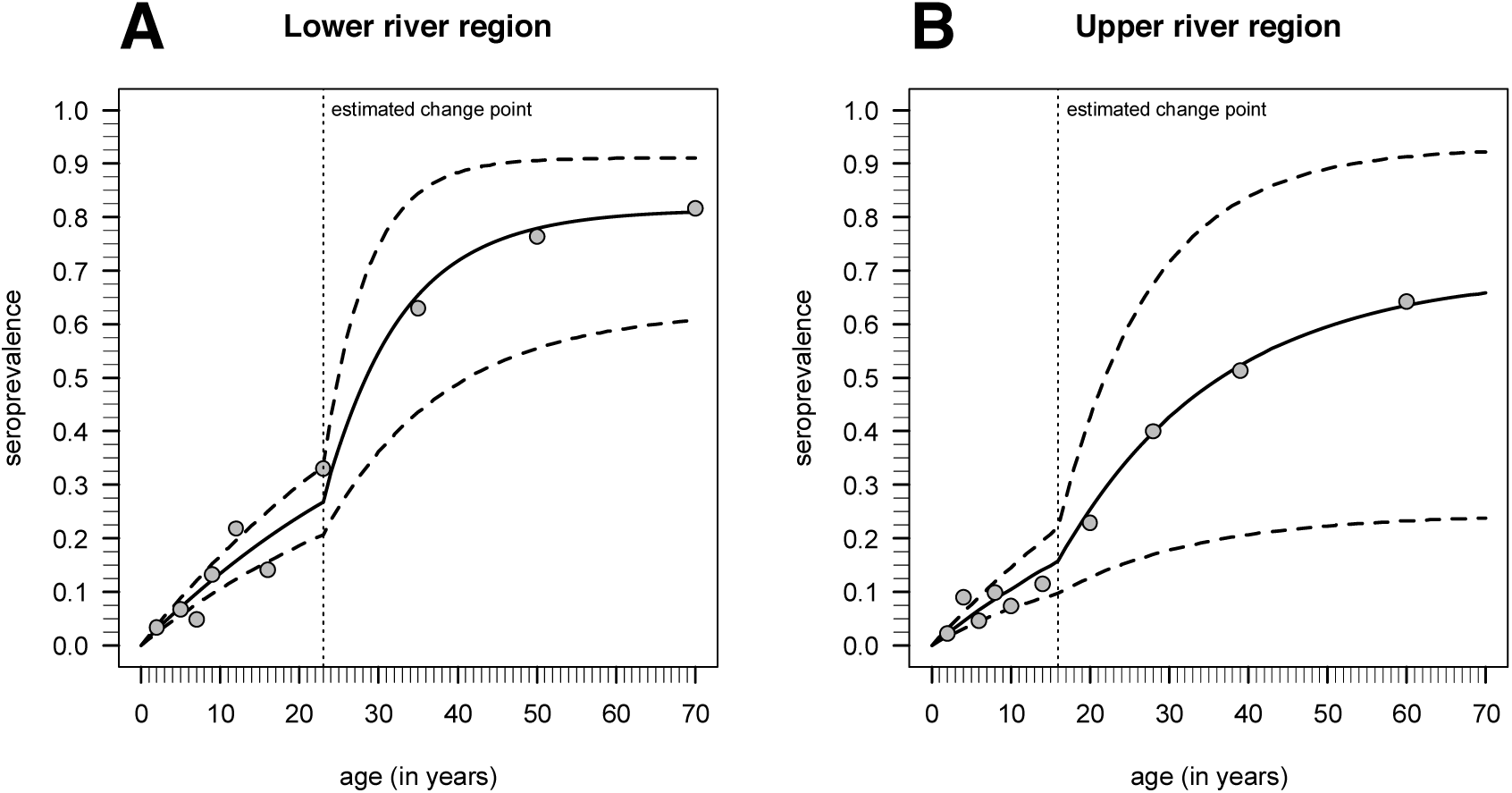
Expected seroprevalence curve as function of age (solid lines) according to the maximum likelihood estimates and the respective 95% confidence intervals (dashed lines) according to the best models selected in Figure 3. The dots represent the observed seroprevalence when the age distribution was broken down in the respective deciles and the vertical pointed line refers to the change point estimated by the profile likelihood method.

A recent serological study using samples from Tanzania^13^ examined the age-specific seroprevalence of anti-Pgp3 antibodies in a trachoma-endemic community that had received two rounds of high coverage azithromycin MDA. The all-ages prevalence of ocular Ct infection had fallen from 9.5% to 0.1% two years after MDA^47^, and to 0% five years after MDA^37^, with a corresponding 11-fold decrease in SCR^13^. This change in infection prevalence occurred in a more defined (and probably narrower) timeframe than the one we have studied in The Gambia. This resulted in a more acute change in SCR, as has been demonstrated in malaria modelling exercises^26,48^. Previous trachoma modelling studies suggest that an individual may require upwards of 100 lifetime ocular Ct infections in order to develop TT^49^, so even a modest reduction in transmission may have significant public health implications and reduce the future incidence of TT.

Research is ongoing to address remaining challenges in interpreting trachoma seroprevalence. It is unclear how many infections are required for seroconversion to occur. Studies involving urogenital Ct infection suggest that just 68% of infected women produce IgG antibodies against Ct^50^. The intensity of the inflammatory response, and the surface area of inflamed mucosa, however, are both likely to differ between the infected conjunctiva and infected female urogenital tract. Additionally, further work is needed to determine the half-life of Pgp3 antibodies and seroreversion rates. A previous study that examined a high-prevalence community before and after one round of azithromycin suggested that individual anti-Pgp3 antibody levels decreased slightly six months after drug treatment, but not enough to be considered seroreversion^51^. This is similar to results seen for urogenital Ct infection, where anti-Pgp3 antibody titres decreased over 4–7 years, but patients were still in the seropositive range^52^. Although we have estimated SRR in this study (Table 5), a more accurate estimate could be obtained from a longitudinal study collecting serum samples over a period of years. In low transmission settings, such as post-MDA communities, the SRR may be underestimated if the assumption of a balance between total number of seropositive and seronegative individuals does not hold true. Further studies to determine SRR are currently underway.

Dichotomising antibody levels to a simple seropositive/seronegative classification provides a straightforward estimation of seroprevalence, but SCR estimates could potentially be improved by using a model based on antibody levels and multiple sampling time points, as suggested by Yman et al^53^. Such models might assume that antibody levels increase with age, as exposure is age-dependent and that transmission intensity can be calculated by measuring the boost in antibody levels. The use of age group-specific geometric mean antibody levels could be explored^53^ in addition to SCR.

Approximately 4% of 1–9-year-olds were positive for antibodies to Pgp3, which may be due to previously acquired ocular Ct infections and/or to ocular or respiratory Ct infections acquired at birth from mothers with urogenital Ct infections^54^. This seroprevalence is within the range of prevalence values previously estimated in post-MDA surveys in Tanzania and Nepal ^12,13^. There was also no observed increase in anti-Pgp3 antibody positivity with age in 1–9-year-olds (Supplementary Table 2), in contrast to what is observed in trachoma-endemic settings, whether treatment-naive ^16^ or after 3 rounds of MDA ^15,19^ and is in stark contrast to communities with a high prevalence of ocular infection^14^. Focusing on age-specific changes in seropositivity as a measure of cumulative exposure to ocular Ct infection might offset antibody responses from peri-natal infection, as the latter would be expected to be consistent across all ages, or even to decline with increasing age. The data from The Gambia presented here, combined with those from a variety of pre-and post-MDA settings, contribute to an understanding of the potential use of antibody-based surveillance of children to ensure a lack of infection recrudescence.

Data from paired pre-and post-MDA surveys, or longitudinal data from MDA surveys, would substantially improve parameters and modeling efforts. A detailed series of surveys in one population would help to develop generalized models for use elsewhere. The inclusion of infection data for both ocular and urogenital Ct infection is needed to further clarify how urogenital Ct contributes to observed seropositivity rates.

## METHODS

### Ethical Review

This study was conducted in accordance with the Declaration of Helsinki. It received approval from the London School of Hygiene & Tropical Medicine Ethics Committee (LSHTM; reference 7059) and The Gambia Government/Medical Research Council Joint Ethics Committee (SCC1408v2). CDC investigators did not engage with study participants.

### Survey methodology

We conducted a population-based, cluster-random-sampled survey was conducted in February-March 2014. The Gambia is divided into geographically-defined census Enumeration Areas (EAs) of approximately 600-800 people each. Sampling by EA is equivalent to sampling settlements with probability proportional to their size^55^. Twenty EAs in each of URR and LRR were randomly selected for participation. Trained field workers sensitised villagers and obtained verbal community-consent from each village chief *(alkalo*). Field workers and *alkalos* compiled a list of households for each EA, from which households were randomly selected for census and recruitment. The study and consent form were explained to the head of each selected household and prospective participants. All members of selected households were invited to participate, regardless of age. Written (thumbprint or signature) consent was obtained from each participant aged ≥18 years, while a parent or guardian provided written consent for each participating child aged under 18 years. Children aged 12-17 years provided assent before participating.

The trachoma graders were experienced in field grading for active trachoma and had received regular training according to PRET^5^ and Global Trachoma Mapping Project (GTMP) protocols^56,57^. After informed consent was received, the grader examined both eyes of the subject using a binocular loupe (2.5×) and a torch. The grader changed gloves between each participant to minimise the risk of carry-over infection. In accordance with the Gambian NEHP policy, antibiotics were provided to individuals with evidence of active trachoma and to residents of their household.

Each participant had a finger-prick blood sample collected onto filter paper (Trop-Bio, Townsville, Australia) using a sterile single-use lancet (BD Microtrainer, Dublin, Ireland). Each filter paper had six extensions, calibrated to absorb 10 µL of blood each. Samples were air-dried for approximately five hours and then placed in individual Whirl-Pak plastic bags (Nasco, Modesto, California) which were stored with desiccant sachets (Whatman, Little Chalfont, UK) at-20°C. All samples were shipped to LSHTM for testing.

### ELISA Assay

Dried blood spots (DBS) were tested for antibodies against Pgp3 according to the method previously described^19^. Briefly, serum was eluted from dried blots spots then applied to a plate coated with Pgp3 protein^15^; known standards were included and assayed in triplicate on each plate. Following incubation, bound antibody was detected with HRP-labelled mouse anti-human IgG(Fc)-HRP (Southern Biotech, Birmingham, USA). Plates were incubated and washed, and then TMB (*KPL, Gaithersburg*, USA) was added to develop the plates. The reaction was stopped with 1N H_2_SO_4_ and optical density was read at 450 nm (OD_450_) on a Spectramax M3 plate reader (Molecular Devices, Wokingham UK). Readings were corrected for background by subtracting the average absorbance of three blank wells containing no serum, using Softmax Pro5 software (Molecular Devices).

### Statistical Methods

Blanked OD_450_ values for samples were normalised against the 200 U standard included on each plate^19^. Laboratory work was undertaken masked to demographic and clinical information. Statistical analyses were carried out using R^58^. Using the “survey” package and assuming a design effect of 2.65^56^, the 95% confidence intervals (CI) were calculated using the Clopper-Pearson interval^59^. Wilcoxon-Mann-Whitney z-scores^60^ were calculated to compare the proportion seropositive between different regions, ages, and genders. The non-parametric test for trend was used to measure the increase in prevalence of anti-Pgp3 antibodies with age.

A finite mixture model^25^ was used to classify the samples as seropositive or seronegative based on normalised OD_450_ values. The data were fitted using maximum likelihood methods, estimating the distribution parameters for each classification group (assumed seropositive or assumed seronegative) as well as the proportion of samples in each category to fit the overall distribution of results^61^. To ensure that the assay had high specificity, the threshold for seropositivity was set using the mean of the Gaussian distribution of the seronegative population plus four standard deviations (the quantile inclusive of 99.994%) of the seronegative population^25,61^.

Population age groups were categorized according to known time points of changes in disease prevalence. The youngest age group was 1–9-year-olds, who in LRR are likely to have been born during or after MDA with azithromycin. The oldest group included people aged 40 years and above, who experienced secular declines in trachoma prevalence prior to 1986, the year of the first National Survey of Blindness and Low Vision^23^. Participants aged between 10 and 39 years were grouped into 10-year categories.

Different reversible catalytic models were applied to the analysis of the data from each region. These models are described as function of seroconversion and seroreversion rates (SCR and SRR, respectively). SCR is defined as the annual mean rate by which seronegative individuals become seropositive upon disease exposure and is usually considered as a proxy of the transmission intensity of the population. SRR describes the annual mean rate by which seropositive individuals revert to a seronegative status in the absence of re-infection. Three reversible catalytic models were fit to the serological data of each region: (i) a simple model that assumes a constant SCR (i.e., transmission intensity) over time and people are born seronegative^27^; (ii) a model with a constant historical SCR until a certain change point in the past where SCR changes abruptly to a new value (current SCR); (iii) a model that also assumes a historical seroconversion rate until a certain change point followed by a time period where SCR decays log-linearly every year. The respective parameter estimation was done using maximum likelihood principles. In particular, the models assuming a change point were estimated via profile likelihood method, as described elsewhere^25^. Model comparison for the data of each region was based on the Akaike’s information criterion where the best model (and change point) is the one that provides the minimum estimate of this statistics.

## ACKNOWLEDGMENTS

We are very grateful to the communities that generously gave their time to participate in this study. We thank the field team and public health staff in The Gambia, Ms PJ Hooper from the International Trachoma Initiative, and Prof Hannah Faal from the University of Calabar and Teaching Hospital. NS is partially supported by the Portuguese Fundação para a Ciência e Tecnologia (Grant # UID/MAT/00006/2013).

The findings and conclusions in this report are those of the authors and do not necessarily represent the official position of the Centers for Disease Control and Prevention.

## AUTHOR CONTRIBUTIONS

S.J.M., R.B., D.C.W.M., A.W.S. and C.h.R designed the study and wrote the paper, D.L.M., G.C., S.G. C.h.R. and S.J.M. developed the protocol for serological analysis, S.J.M. performed serological analysis, S.J.M., C.h.R and N.S. performed the statistical analysis, R.B. trained the trachoma graders, S.J.M., H.J., P.M., R.B., S.E.B., D.C.W.M. and C.h.R. organized field data collection. H.J. and P.M. performed the field grading of trachoma. H.P. provided critical feedback on the paper. All authors reviewed and approved the manuscript.

